# Derivation of cardiac reference ranges for *Mus musculus* using the Collaborative Cross and identification of new cardiac models

**DOI:** 10.1101/2025.01.08.631983

**Authors:** Julia L. Popp, Krishna Patel, Danila Cuomo, Rachel M. Lynch, Steve W. Leung, Brian R. Berridge, Ivan Rusyn, Weihsueh A. Chiu, David W. Threadgill

## Abstract

Contemporary approaches for developing interventions and assessing pre-clinical cardiovascular risk frequently utilize animal and *in vitro* models. However, these models currently lack normal species-specific reference ranges similar to what exists for humans. The genetically diverse Collaborative Cross (CC) population that models human genetic heterogenetiety was characterized to develop mouse-specific cardiac reference ranges for *Mus muscuslus*, the most commonly used pre-clincial model. Heart function was analyzed in males and females from 58 CC strains and C57BL/6J using high-frequency ultrasound under both conscious and anesthetized conditions, as well as conscious electrocardiography to develop two standard deviation-based reference ranges. The sources and magnitude of measurement variability were identified, and inter-laboratory comparisons determined to quantify phenotypic robustness and heritability. Strain was the largest source of variability, while laboratory where data were collected was also significant but sex was not. Additionally, strains were identified that have characteristics of disease-associated phenotypes in cardiac function and electrophysiology similar to human cases including dilated cardiomyopathy, systolic cardiomyopathy, cancer therapy-related cardiac dysfunction, and long QT. These new models allow a more natural, and therefore more translatable progession to a cardiac disease state, supporting development of strain-specific models for cardiac pathologies, ultimately allowing more accurate diagnoses and informative safety assessments in humans.

**Article Summary:** The Collaborative Cross (CC) mouse reference population, modeling human genetic diversity, was utilized to derive cardiac reference values for *Mus musculus*. Strain was the primary source of variability, with the laboratory also playing a significant role, while sex had no impact. Despite inter-laboratory reproducibility challenges, the study identified strains resembling disease-associated phenotypes that offer a more natural progression to cardiac disease and provide crucial data for early disease identification and accurate safety assessments in humans.

## Introduction

Heart disease is the leading cause of death worldwide (WHO 2020), with over 650,000 Americans dying of heart disease each year and accounting for 1 in 4 deaths (VIRANI *et al*. 2020). Although the underlying causes of heart disease are multifactorial, abnormal cardiac phenotypes such as long QT, increased heart size (cardiomyopathy), and low ejection fraction (EF) are important manifestations of cardiovascular disease. Susceptibility to these abnormalities are heritable, with additional exogenous factors such as diet, exercise, metabolic disorders, environmental exposures, and drugs further increasing risk for adverse cardiac events (VIRANI *et al*. 2020). For example, abnormalities in left ventricular (LV) structure and function are heritable (POST *et al*. 1997; JIN *et al*. 2011), and it has become increasingly apparent that LV function is key to heart failure (PONIKOWSKI *et al*. 2016). With highly limited regenerative capabilities and many maladaptive processes, the heart is an especially vulnerable organ to the effect of exogenous factors. Indicative of this vunerability, 33% of all drugs withdrawn from the market between 1998 and 2008 were due to cardiac complications (MACDONALD AND ROBERTSON 2009).

Contemporary *in vivo* approaches for developing new interventions and for cardio-safety evaluation include assessments for cardiovascular structure and function, frequently performed in a variety of animal and *in vitro* models (BURNETT *et al*. 2021), but rarely include considerations of genetic background as a potential contributing risk factor. Of various animal models, non-human primates, dog, and swine are typically preferred pre-clinical species; however, they are not population-based or genetically-tractable models. By contrast, mice differ in heart physiology and electrophysiology from humans and are typically not used in pre-clinical safety studies, but have the advantage of the genetic modification tools and reference populations to study genetic causes of human disease. Even though mice are used to study cardiac pathology (BERRIDGE *et al*. 2016), no standard cardiac reference ranges exist for *Mus musculus* similar to that in humans. Characterization of the population diversity in cardiac phenotypes in mice is important for identifying individuals at increased risk for adverse cardiac events and for using mice in pre-clinical safety testing.

Previous reports, such as those derived from the Framingham Heart study (WILD *et al*. 2017; ANDERSSON *et al*. 2019), have provided a wealth of information to determine normal variation in baseline cardiac physiology in humans (FOX *et al*. 2013; AUNG *et al*. 2019). Human data is presented as reference ranges, which are typical values within two standard deviations from the population mean. They are used to describe normal ranges of cardiac phenotypes in people and are often based on sex and age. Although previous studies have reported variable outcomes in cardiovascular disease states based on genetic factors in humans and animal models (MARON AND MULVIHILL 1986; GOLBUS *et al*. 2012; SALIMOVA *et al*. 2019), reference ranges have not been derived for animal models that can be used to evaluate baseline cardiac risks before intervention trials. For example, the C57BL/6 mouse is one of the most commonly used strains in experimental research (SIMON *et al*. 2013), and has been used to model cardiovascular therapies for humans (LEE *et al*. 2019; MCLAUGHLIN *et al*. 2019). Yet, the hearts of C57BL/6 mice are deemed as a model for hypertrophic “athlete’s heart” that is not representative of most people (HOIT *et al*. 2002).

The role of genetic variability in disease risk is best modeled in the mouse, which has the most extensive genetic resources including dozens of strains, each potentially representing a unique risk status for heart disease development. Furthermore, nearly all environmental factors can be controlled in mouse studies, supporting detailed phenotypic and physiologic analyses. However, before the mouse can be fully exploited as a model of human population variation in cardiac physiology and disease, detailed baseline characterizations across strains and sources of measurement variation are required to properly identify models representing various risk statuses for translation to humans. To overcome the limited combinations of genetic variation present in mouse strains, the Collaborative Cross (CC) was developed as a mouse genetic reference population that has sufficient diversity to model the range of genetic diversity present in the human population (THREADGILL, HUNTER AND WILLIAMS 2002; CHURCHILL *et al*. 2004; ROBERTS *et al*. 2007).

To better understand population-based variation in cardiac phenotypes that contribute to differential disease susceptibility, and to provide a model for human population variation, this study reports detailed characterization of strain-dependent differences in baseline cardiovascular physiology, species-specific reference ranges, and sources of variation using echocardiography and electrocardiography (ECG) based on the CC population. Echocardiography is used as a fast and reliable assessment of cardiac morphology and function. However, there is controversy as to whether measures on conscious or sedated mice are more reliable and representative of baseline cardiac function because stress of handling and the possibility of anesthesia inducing changes in cardiac function (PACHON *et al*. 2015). To compare the approaches, both conscious and isoflurane-sedated echocardiography were performed in both sexes of 58 CC mouse strains and C57BL/6J as a reference control to evaluate strain-dependent cardiac physiology. Conscious ECG was also performed to evaluate the electrical activity and autonomic regulation of heart function. Early contractile dysfunction can be detected by subtle changes in echocardiography by global longitudinal strain (GLS), EF, and LV mass, and in ECG by prolonged repolarization and depolarization of the myocardium during various intervals of the cardiac cycle, which are dependent on baseline heart function. The measures were then used to derive reference ranges for normal cardiac physiology in *Mus musculus* and to identify new models of predispotion to human cardiac conditions. Given the complexity of cardiac physiology and disease, the use of the CC will aid in elucidating genetic and associated phenotypic networks to support identification of genes underlying risk and poorly adaptive phenotypes, particularly in response to exogenous changes such as disease, toxicants, and diet in a population-based context.

## Materials and methods

### Mice

Three male and three female adult mice, 6-9 weeks of age, from 25 CC strains were bred at the University of North Carolina Systems Genetics Core (UNC), and 33 CC strains and C57BL/6J strain were from colonies bred in house that originated from UNC for a total of 354 mice. Mice were provided diet 2919 (Envigo) except for CC004/TauUnc, CC007/Unc, CC008/GeniUncJ, CC009/UncJ, CC010/GeniUncJ, CC018/UncJ, CC024/GeniUncJ, CC026/GeniUnc, CC028/GeniUncJ, CC031/GeniUnc, CC033/GeniUncJ, CC038/GeniUnc, CC039/Unc, CC040/TauUnc, CC042/GeniUnc, CC045/GeniUnc, CC051/TauUnc, CC059/TauUnc, CC060/UncJ, CC068/TauUncJ, CC071/TauUnc, CC074/Unc, CC075/UncJ, CC079/TauUncJ, CC080/TauUncJ, CC081/Unc, and CC084/TauJUnc, which were fed 8604 (Envigo) and water *ad libidum* using a 12-h light/dark cycle. These diets that differ by only 5% in fat composition were not found to impact cardiac measures during analysis. Housing and experimentation rooms are kept constant at 21°C-23°C. Mice were housed in cages of 2-5 mice, with clean bedding and nestlets. Strains will subsequently be referred to without the laboratory code for brevity. On Day (D) 1 and D2, mice were subjected to conscious ECG with D3 being a rest day for the mice. Anesthetized echocardiography and body weights was obtained on D4, and on D5 mice were subjected to conscious echocardiography (Fig. 1). Since mice were used for further unrelated studies, no humane endpoints were necessary. No criteria were set for exclusion of any animals.The investigation conforms with the 2011 Guide for Care and Use of Laboratory Animals published by the National Institutes of Health. All experimental protocols were approved by the Institutional Animal Care and Use Committee (IACUC) of Texas A&M University (IACUC No: 2019-0435).

**Fig 1.**
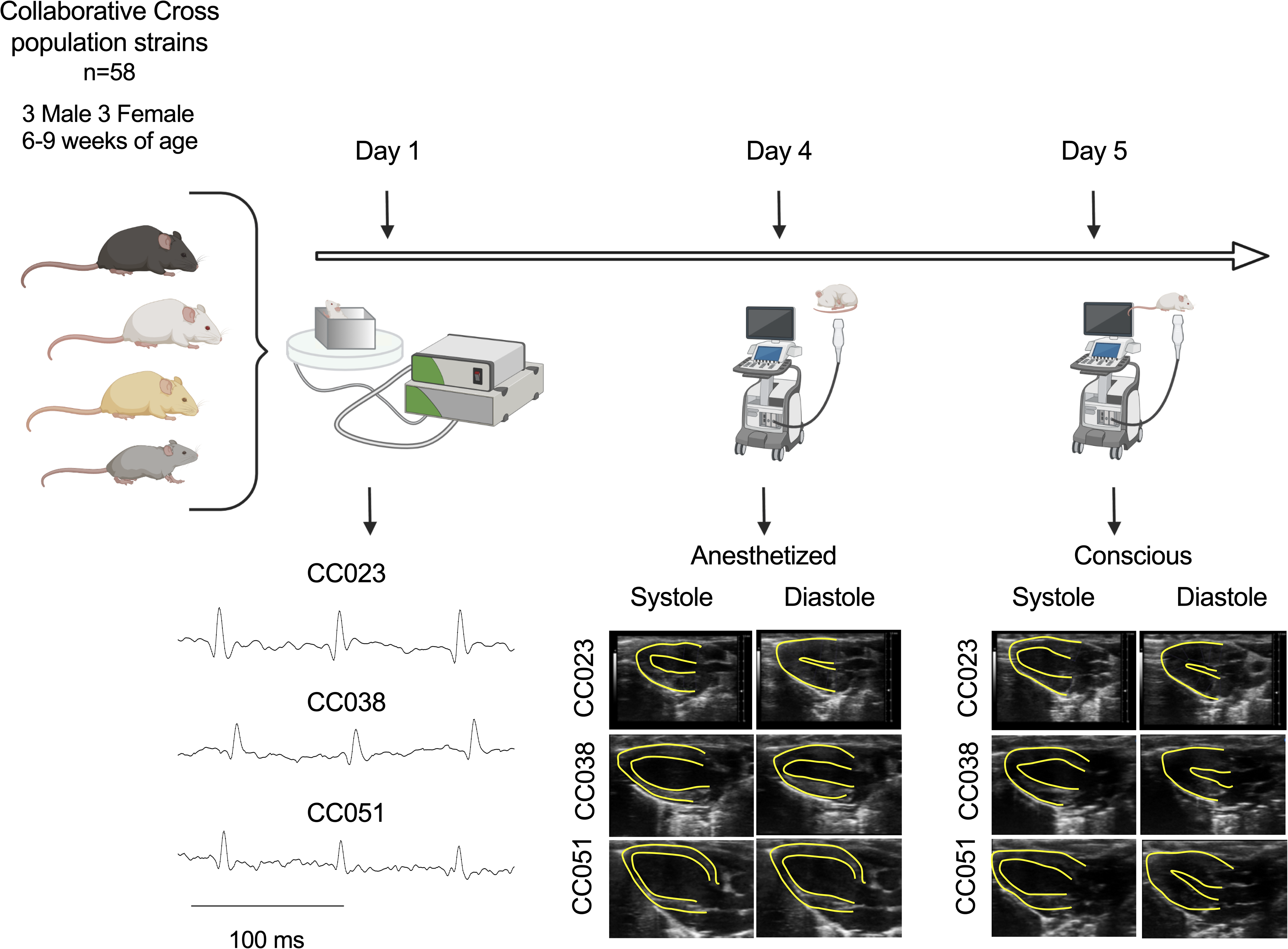
Experimental Design. Mice from 58 CC Strains and C57BL/6 were subjected to ECG (Day 1 and 2), anesethetized echocardiography (Day 4), and conscious echocardiography (Day 5).

### Electrocardiograhpy

ECG recordings were obtained non-invasively using an ECGenie apparatus (Mouse Specifics). In order to eliminate circadian influences, ECG was performed between 9 am and 12 pm. Briefly, individual mice were removed from their home cage and placed on the neonate recording platform 2 minutes prior to data collection to allow for acclimation. ECG signals were acquired through disposable footpad electrodes located in the floor of the recording platform. The amplitude of the ECG signal depends on the passive nature of detecting the signals through the mouse paws that is influenced by experimental variables such as how the mouse stands, the weight of the animal, how much pressure is exerted on each paw, whether there is any moisture on the lead plate (*i.e*. urine and feces), and the age of the lead plate (oxidation). Consequently, amplitude differences were not used in the analysis. Approximately 20-40 raw ECG signals were analyzed per mouse using e-MOUSE software (Mouse Specifics), which employs processing algorithms for peak detection, digital filtering, and correction of baseline for motion artifacts. Heart rate (HR) was determined from R-R intervals, and HR variability was calculated as the mean difference in sequential HRs for the entire set of ECG signals analyzed. The software also determined cardiac intervals (*i.e.,* PR, QRS, and QTc) with rejection of spurious data resulting from noise or motion. In mice, the T-wave often merges with the final part of the QRS complex (WEHRENS, KIRCHHOFF AND DOEVENDANS 2000), so the end of the T-wave of each signal was automatically defined as the point where the signal intersects the isoelectric line. All data were collected by a single operator (JLP).

### Transthoracic echocardiography

Cardiac morphological and functional parameters were assessed with a Vevo3100 high-frequency ultrasound imaging system (FUJIFILM VisualSonics). For anesthetized echocardiographic analysis, mice were anesthetized with 4% isoflurane in oxygen and placed in a supine position on a temperature-controlled 39°C imaging platform under continuous supply of 1.3%-1.8% isoflurane through the mouth and nose. Ventral hair on the rostral thoracic cavity was removed at least one day prior using a hair removal cream (Nair), and pre-warmed ultrasound transmission gel was applied on the chest at the image location. Body weights were taken after anesthetized transthoracic echocardiography. Mice were allowed to rest for 24 hours before conscious echocardiography. For conscious echocardiographic analysis, mice were held firmly in the palm of one hand by the nape of the neck in the supine position with the tail held between the last two fingers. Pre-warmed ultrasound gel was placed on the chest at the image location. For two-dimensional (2D) imaging (B-mode) view along the parasternal long axis, the transducer was placed vertically to the animal body on the left side of its sternum with the notch of transducer pointing to the animal’s head. After an imaging session, ultrasound gel was removed with water dampened gauze. In order to eliminate circadian influences, ultrasounds was performed between 9 am and 11 am. Measurements and calculations were performed using VevoLAB strain analysis (v3.2.6) software package (FUJIFILM VisualSonics) on three consecutive beats according to the American Society for Echocardiography (MOR-AVI *et al*. 2011). Papillary muscles were excluded from the cavity in the tracing. EF was calculated as EF% = [(EDV− ESV)/EDV] x 100 (End diastolic volume = EDV, end systolic volume = ESV). GLS is a measure of deformation of the myocardium on the long axis of the heart and was calculated as [(LV Length – LV Length_0_)/LV Length_0_]; a smaller negative value reflects a more impaired GLS. Fractional shortening (FS) was calculated as FS% = [(LVIDd − LVIDs)/LVIDd] x 100 (LV internal diameter at diastole = LVIDd, LV internal diameter at systole = LVIDs). The stroke volume (SV) was calculated as EDV-ESV. Cardiac output (CO) is calculated as SV*HR. HR was determined from the cardiac cycles recorded on the M-mode tracing using at least three consecutive systolic intervals. Results are provided as reference ranges and mean ± SD of the mean for 6 mice from each strain (3 mice from each sex). All data were collected by the same operators (JLP and subsets were confirmed by a second operator KP).

### Statistical analysis

GLS negative values were converted to positive. Shapiro-Wilk statistic was used for normality on untransformed and log-transformed values; higher p-value was the preferred transformation. For characterizing sources of variance in this study, ANOVA was performed for each variable with factors strain, batch (nested in strain), sex, and body weight. For cross-lab comparisons, laboratory (nested in strain) was added as a factor, but because only male mice with no batch information were available for other studies, batch and sex were not included. The ω^2^ statistic was used as the measure of “effect size” for a factor, *i.e*., the fraction of variance the factor explains (OLEJNIK AND ALGINA 2003). The overall and strain-specific variances are also calculated, as is the residual variance representing intra-strain variation. Computations were performed using R studio version 1.4.1106, with ω^2^ computed using the sjstats package (version 0.18.1). Heritabilities were estimated using H^2^ = σ^2^_G_/( σ^2^_G_ + σ^2^_E_) with σ^2^ being the standard deviations of the means from all lines (across lines) and σ^2^_E_ being the mean of the standard deviations of individual lines (within lines).

## Results

### Variation in heart morphology and function across CC strains under anesthesia

Echocardiography under anesthesia is the standard approach for analyzing heart morphology and function in experimental models. Inter-strain variation was found in heart size-dependent phenotypes like EDV (35.64 μL ± 11.53), ESV (20.02 μL ± 9.05), diastolic LV mass (65.99 mg ± 13.84), and systolic LV mass (64.42 mg ± 13.88) (Table 1). Derived phenotypes, calculated from the primary phenotypes such as mass and volume, combine systolic and diastolic values to generate functional values of cardiac physiology. These include SV (15.61 μL ± 5.74), FS (22.73% ± 9.57), CO (6.27 mL/min ± 2.78), EF (44.85% ± 13.67), and GLS (-14.99% ± 5.96). Strain, sex, body weight, and batch within strain were analyzed for their potential contribution to measurement variation observed in the CC population. Mouse strain, or genetic background, was a significant contributor to variability (p < 0.05) in HR, EF, and GLS (Fig. 2). While strain was the largest defined contributor to variability in HR and EF, residual factors also contributed to each of the three clinically relevant phenotypes. Heritability of the variation in HR, EF, and GLS were found to be high (H^2^ = 0.50, 0.48, and 0.57, respectively). No significant sex effects were found.

**Fig 2.**
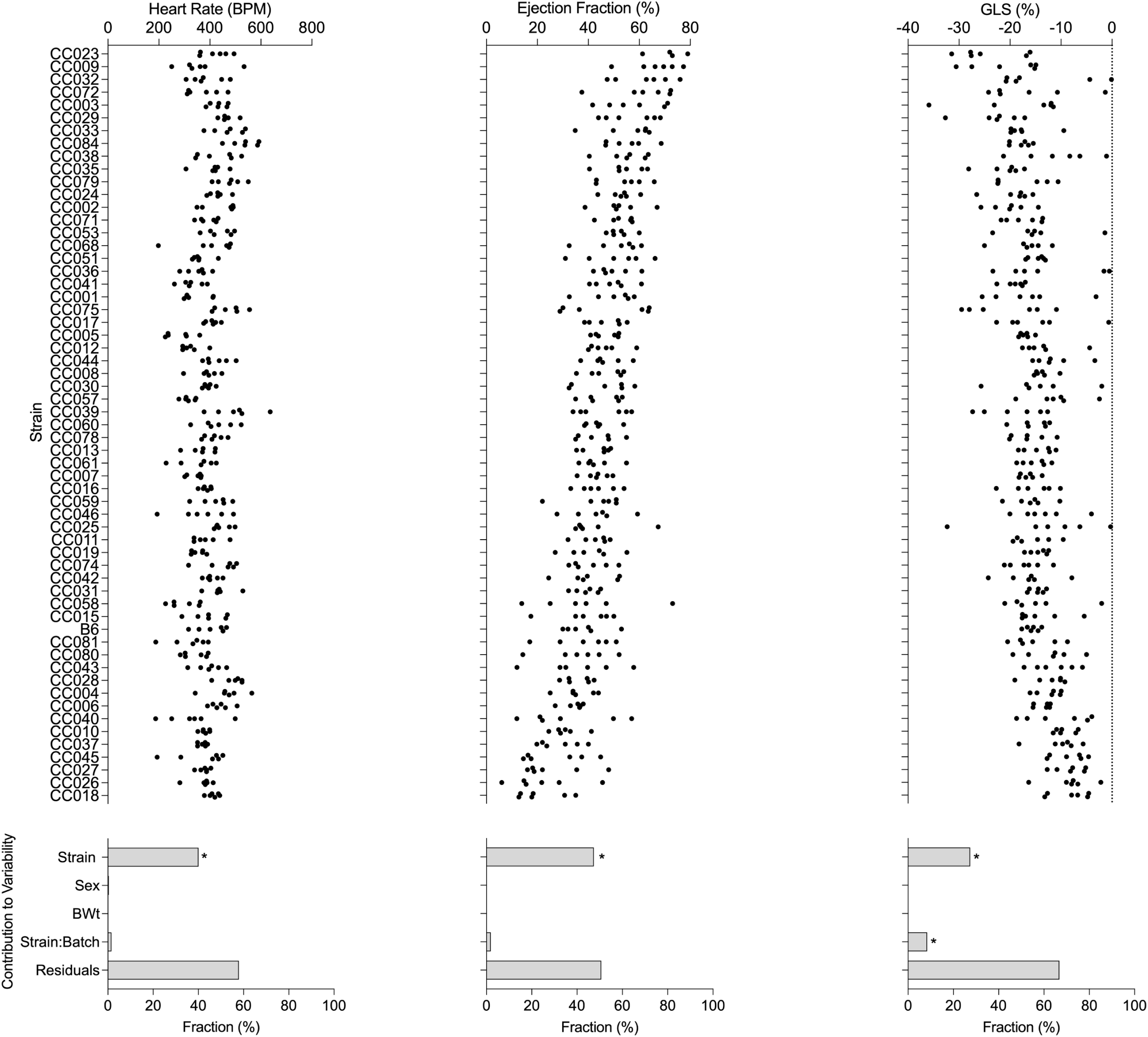
Anesthetized echocardiography elicits inter- and intra-strain variability in cardiac phenotypes. HR, EF, and GLS demonstrate variable response to anesthesia based on strain. Strain is a contributing factor to the differential response, while sex and body weight were not.

**Table 1.**
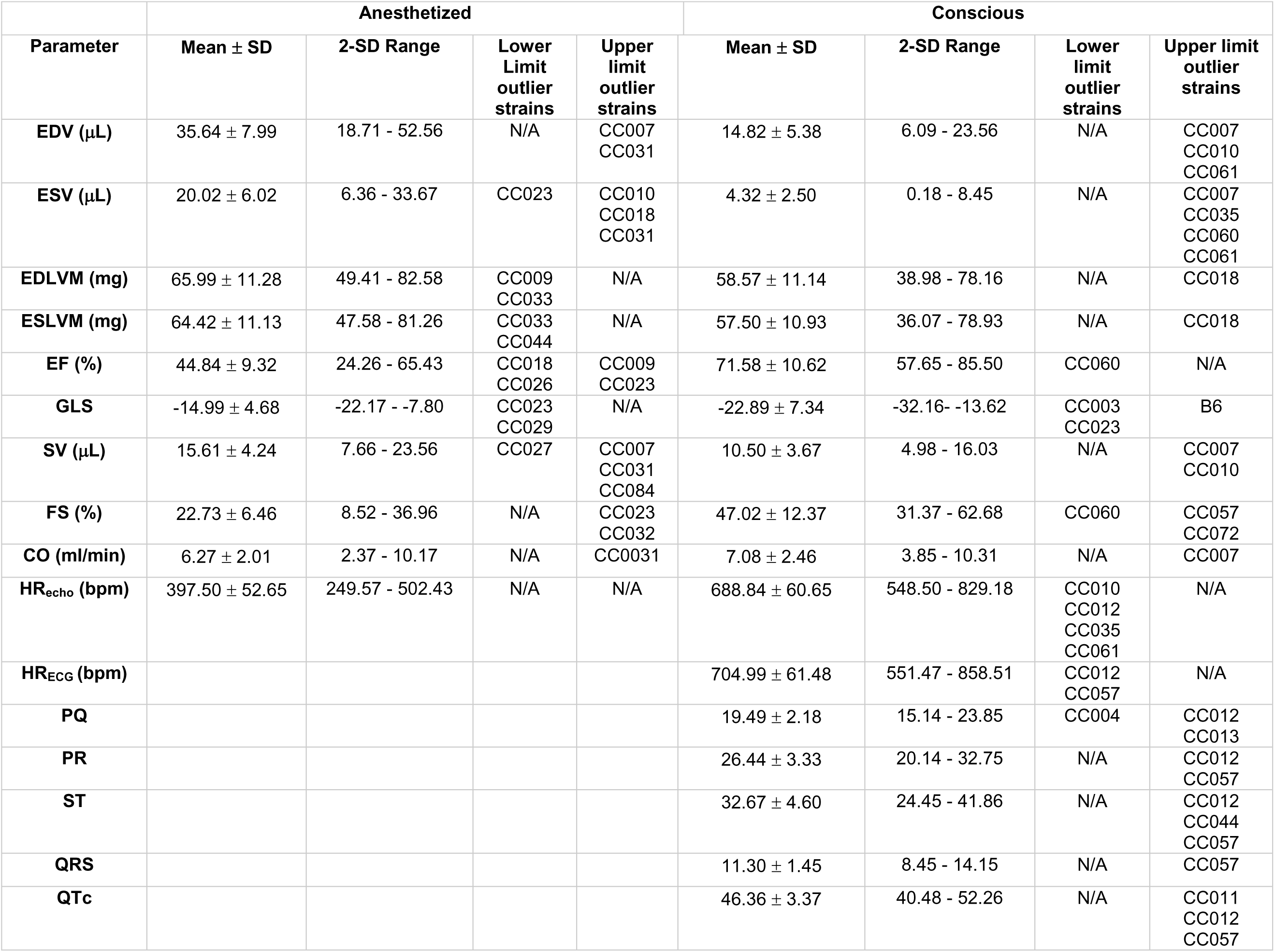
Reference ranges for Mus musculus and outlier strains.

### Differential response to anesthetized and conscious echocardiography in CC strains

The use of anesthetized mice rather than conscious mice for echocardiography has been debated in the research community (ROTH *et al*. 2002). To determine the impact of anesthesia on echocardiography, the same animals were analyzed under conscious and anesthetized conditions. For physiologic HR of conscious mice at rest, ranges between 450-500 beats per minute (bpm) were reported in BALB/c mice using telemetry electrocardiography (KRAMER *et al*. 1993), and 658 bpm ± 9 in 129S6/SvEvTac mice using conscious echocardiography (YANG *et al*. 1999). Studies using anesthetized echocardiography generally keep HR in a predefined range to capture the corresponding ideal heart function as determined by EF, but in a genetically diverse population such as the CC, HR is correlated with EF under anesthetized but not conscious echocardiography (Spearmans p = 0.0010 and 0.7839, respectively).

Across all CC strains under conscious echocardiography, HR was 688.84 bpm ± 100.39; EDV (14.82 μL ± 7.01) and ESV (4.32 μL ± 3.52); LV mass at diastole (58.57 mg ± 15.11) and systole (57.50 mg ± 15.64); and SV (10.50 μL ± 4.52), FS (47.03% ± 14.39), and CO (7.08 mL/min ± 2.86). EF (71.58% ± 12.53) and GLS (-22.89% ± 8.52) also showed differential response to conscious echocardiography across CC strains. Using an inbred strain or outbred stocks, it has been reported that anesthetics cause strain-dependent effects *in vivo* (SONNER, GONG AND EGER 2000), depresses cardiac function (CHAVES, WEINSTEIN AND BAUER 2001; ROTH *et al*. 2002), and that sympathetic tone exerts the predominant influence on HR in conscious mice (JUST, FAULHABER AND EHMKE 2000). Effects on sympathetic tone was observed in the current study when comparing both methods of echocardiography. Conscious intrastrain HR variability increases as HR values decrease, indicating that sympathetic tone is strain-dependent, and at lower, more relaxed HRs, higher variability indicates more influence from other factors. Strain was a significant contributer to variability (p < 0.05) in the changes (Δ) of HR, EF, and GLS between anesthetized and conscious echocardiography (Fig. 3). Heritability in conscious HR, EF, and GLS was 0.46, 0.60, and 0.61, respectively. Additionally, the heritability of the delta of these phenotypes between conscious and anesthetized echocardiography was high at 0.53, 0.55, and 0.59, respectively.

**Fig 3.**
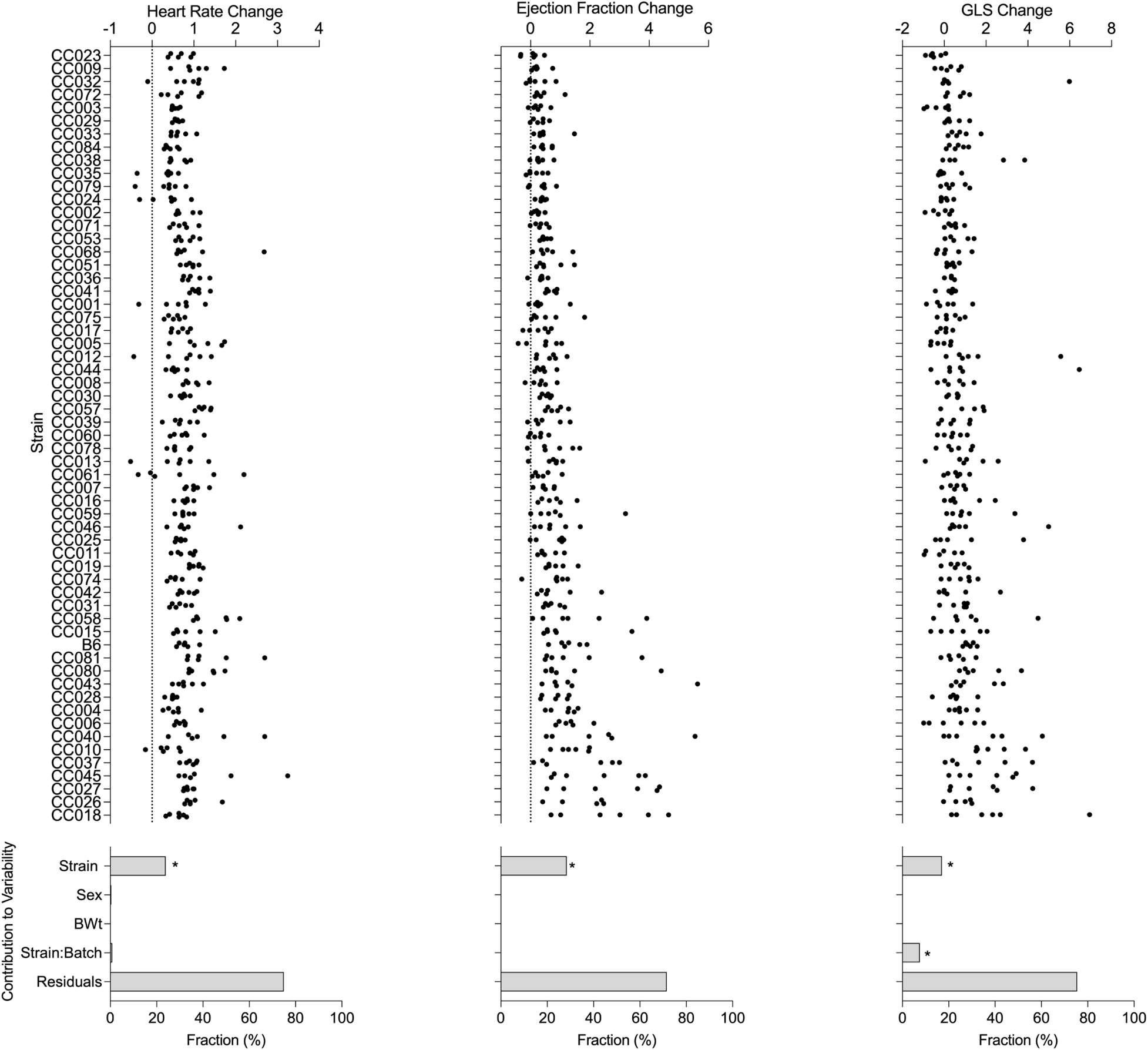
The delta between conscious and anesthetized echocardiography elicits inter- and intra-strain variability in cardiac phenotypes. HR, EF, and GLS demonstrate variable response to anesthesia based on strain. Strain is a contributing factor to the differential response, while batch, sex, and body weight were not.

### Variation in heart electrophysiology across CC strains

Conscious ECG revealed considerable differences across strains (Fig. 4). HR was 712.58 bpm ± 60.30. PQ (19.49 ms ± 2.48) and PR (26.54 ms ± 3.33) interval duration represent intra-atrial conduction and excitation delay within the atrioventricular (AV) node. Prolongation of these intervals indicates AV block and is predictive of adverse outcome in humans (CHENG *et al*. 2009; RASMUSSEN *et al*. 2017). The ST interval (32.66 ms ± 4.60) pathologies include elevation, which can lead to acute myocardial infarction. The QRS complex interval (11.30 ms ± 1.45) showed wide variation across strains, and represents ventricular depolarization, and a prolongation (“wide QRS”) usually indicates abnormalities in the interventricular septum due to underlying disease (BRENYO AND ZAREBA 2011). QTc is a HR-corrected QT interval, whose prolongation can lead to *torsades de pointes* or sudden cardiac death. QTc intervals (46.36 ms ± 3.34) were also variable. Strain was a significant contributor to variability (p < 0.05). While strain was the highest characterized contributor to variability, residual factors contributed to each of HR, PR, QRS, and QTc. Heritability of HR, PR, QRS, and QTc was 0.44, 0.51, 0.50, and 0.53, respectively.

**Fig 4.**
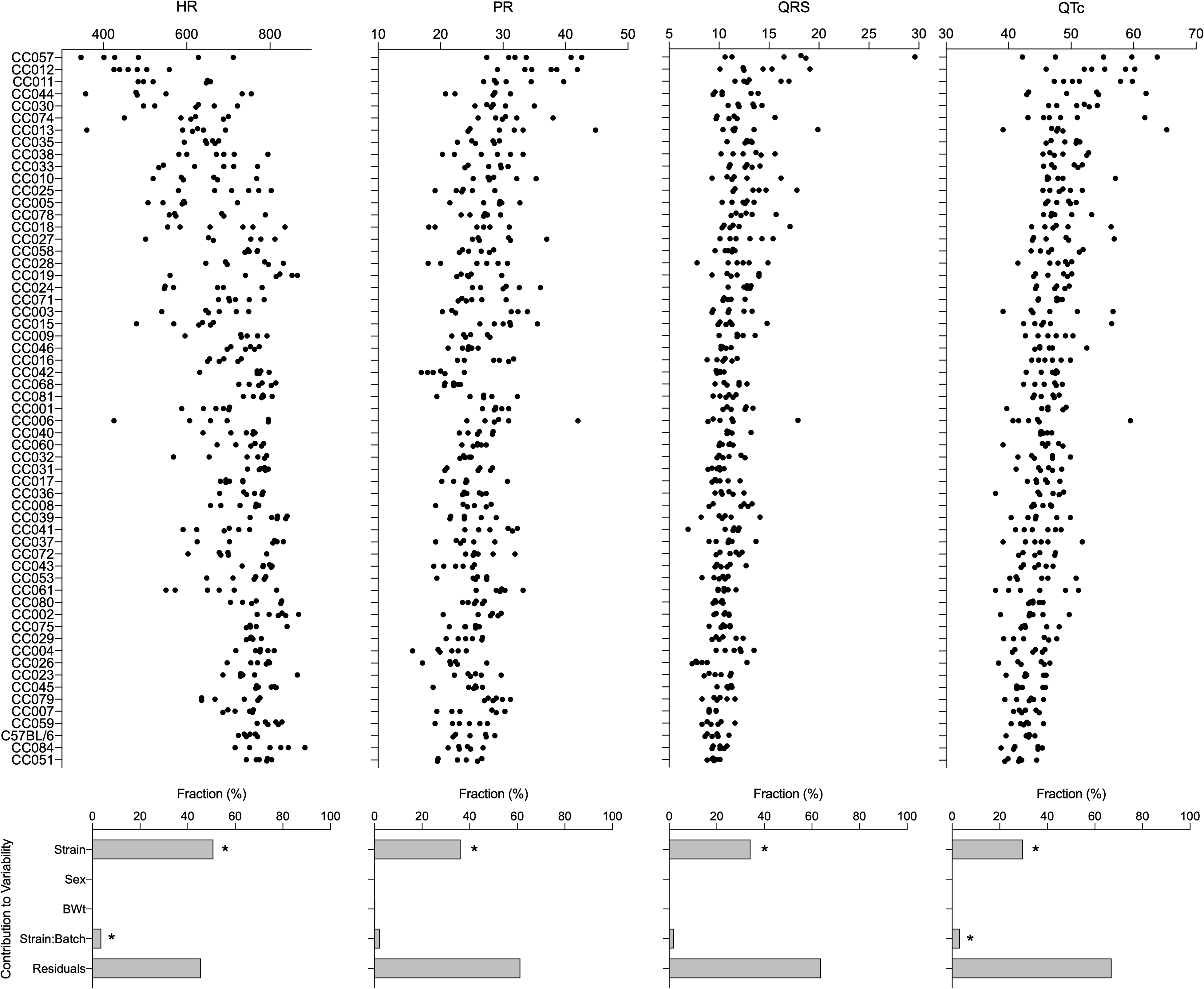
Concious ECG elicits inter- and intra-strain variability in cardiac phenotypes. HR, PR, QRS, and QTc demonstrate variable response to anesthesia based on strain. Strain is a contributing factor to the differential response, while sex and body weight were not.

### Reproducibility of CC mouse cardiac phenotypes

The CC are genetically reproducible and ideally, phenotypes collected under similar conditions should be comparable between laboratories. To evaluate cardiac phenotype reproducibility, the baseline data collected in this study were compared with previous studies that used a subset of CC strains. Salimova et al. assessed parameters of cardiac morphology and function using anesthetized echocardiography before and after myocardial infarction (SALIMOVA *et al*. 2019). Their baseline measurements were collected at 12 weeks of age, one month older than in the current study, and revealed marked strain-dependent differences in body weight, LV mass, EF, and SV. A total of 15 strains (14 CC and one founder strain, only male mice) overlapped with the current study. The previous study performed echocardiography under 1.5% isoflurane with HRs of most strains remaining between 400 and 450 bpm. Similarly, the current study kept isoflurane anesthesia at 1.5% except for a few strains where it ranged between 1.3% and 1.8% due to differential requirements of anesthesia to maintain sedated state. Data sets were combined for analysis and evaluated for sources of variability. Even after combining data across laboratories, strain remained a significant contributor to variability (p < 0.05) for LV mass, EF, EDV, and ESV. Body weight was a significant contributor to variability (p < 0.05) since LV mass, EDV, and ESV are dependent on body weight. Laboratory was also a significant factor in all four common phenotypes (p < 0.05), but to a lesser extent than strain. Heritability of Salimova et. al data for LV Mass, EF, EDV, and ESV was 0.28, 0.48, 0.45, and 0.56, respectively, which compares well with the current study with 0.57, 0.48, 0.49, and 0.47, respectively (Fig. 5).

**Fig 5.**
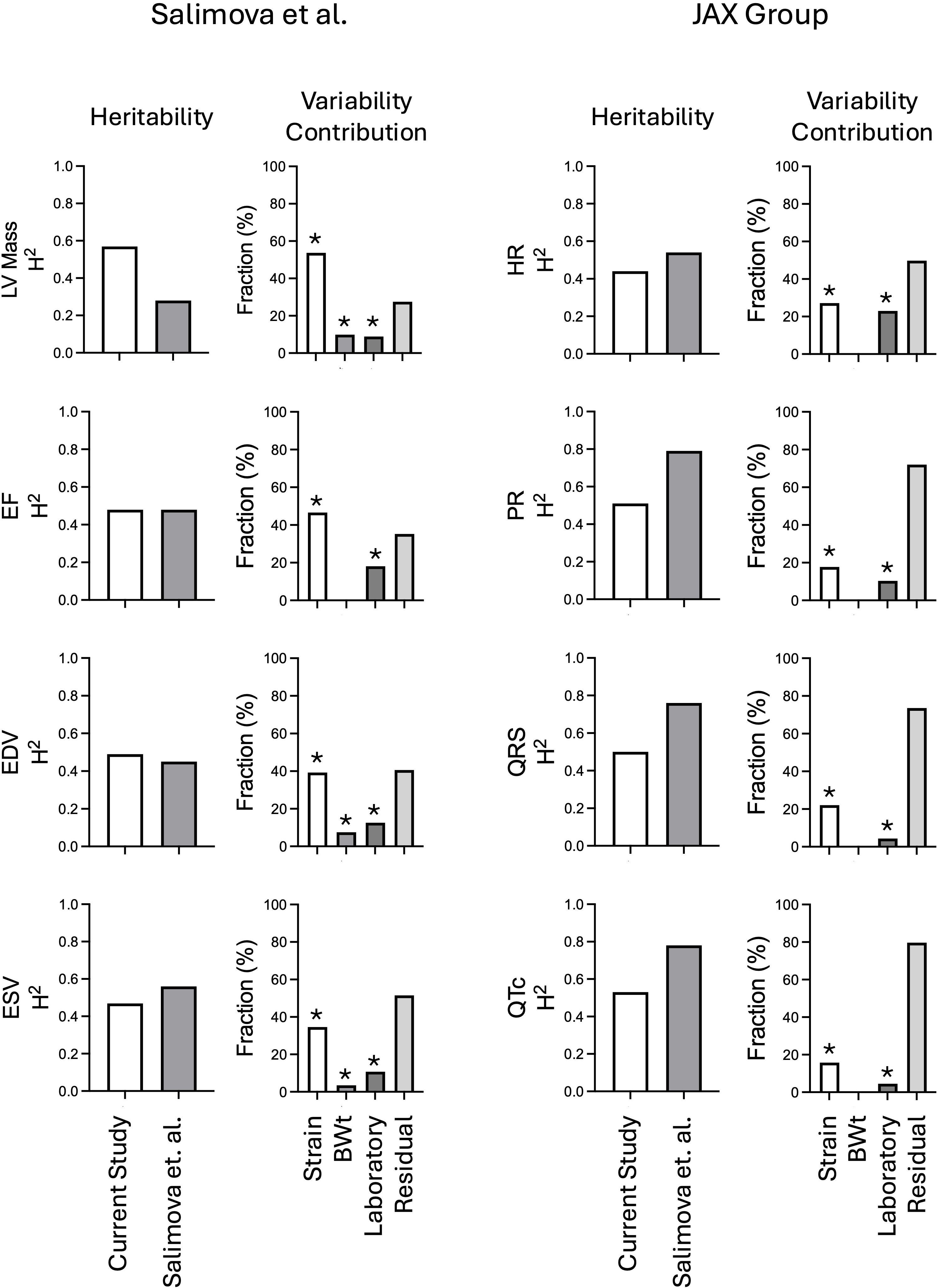
Comparison of data sets between laboratories reveals variability to reproducibility. Baseline cardiac phenotype data collected in this study was compared with previous studies that used a subset of CC strains. Salimova et al. assessed parameters of cardiac morphology and function using anesthetized echocardiography. Jax Lab performed conscious electrocardiography on another subset of strains present in this study. Heritability (H^2^) and assessment of variability contribution by population characteristics.

The Jackson Laboratory (Jax) also reported conscious electrocardiography on 18 CC strains of both sexes at 10-12 weeks of age (LABORATORY 2019). In those studies, data were collected using an open, elevated recording platform rather than a small neonate box, and data were collected during the first 5-10 minutes, as compared to waiting 2 minutes for the current study. Data sets were combined for analysis and evaluated for sources of variability. Strain was a significant contributer to variability (p < 0.05) for HR, PR interval, QRS interval, and QTc. Laboratory or measurement instrument was also a significant factor in three of the common phenotypes (p < 0.05), but to a lesser extent than strain. The contribution of laboratory to HR variability is much higher than for other phenotypes, consistent with the hypothesis that HR is highly dependent on environment. In the Jax data, heritability for HR, PR interval, QRS interval, and QTc was 0.54, 0.79, 0.76, and 0.78, respectively, while in the current study it was 0.44, 0.51, 0.50, and 0.53, respectively.

### Cardiac reference ranges for *Mus musculus*

There are no established reference range values for *Mus musculus* cardiac measurements similar to what exists for humans and other pre-clinical species. To develop mouse-specific values, data across CC strains were used to develop reference ranges for cardiac measures in mice (Table 1). There were no sex-specific differences, so male and female data were combined to n = 6 mice for each strain. Because human reference ranges exlude patients with outlying values of blood pressure, glomerular filtration rate, and cholesteral, and history of hypertension and diabetes, strains that fall outside of the population reference range are excluded for the normal mouse reference range. Reference ranges were calculated to be 2 standard deviations on either side of the population mean for each phenotype. Coefficient of variation (CV) of normal mouse means were also compared with available official human data from the American Society of echocardiography (LANG *et al*. 2015).

### Anesthetized echocardiography cardiac phenotypes

In anesthetized mice, average HR is 397.50 ± 52.65 bpm, and the HR normal reference range of anesthetized mice is 249.57 bpm – 502.43 bpm, with all strains being within the reference range as would be expected since anesthesia kept heart rates within narrow ranges. EF normal reference range under isoflurane 1.5% anesthesia is 24.26% - 65.43%, wider than the 50% to 65% previously reported using individual inbred strains or outbred populations lacking the diversity present in the CC (YANG *et al*. 1999; STYPMANN *et al*. 2006). CC009 and CC023 are above the population range at 66.2% and 74.7%, respectively, while CC018 and CC026 are below the reference range at 21.3% and 22.0%, respectively. GLS normal reference range under the same conditions was -22.17% to -7.80% with CC023 and CC029 being below the reference range at - 24.3% and -23.0%, respectively.

### Conscious echocardiography phenotypes

The normal reference range for conscious HR is 548.50 – 829.18 bpm with numerous strains, including CC010 (lowest at 505 bpm), CC012, CC035, and CC061, being below the lower limit and none above. For conscious EF, the normal reference range is 57.65% - 85.5%, slightly wider than the 65% - 84% previously reported using individual inbred strains or outbred populations (YANG *et al*. 1999; POPOVIC *et al*. 2005). Only CC060 was below the reference range at 53.4%. GLS normal reference range is -32.16% to -13.62 with CC003 and CC023 being above at -12.9% and -13.4%, respectively, and B6 being below at -34.3%.

### Electrophysiology phenotypes

The normal reference range for HR during conscious ECG is 551.47 – 858.51 bpm, which is very similar to the reference range determined by unconscious echocardiography. CC012 and CC057 had HR of 478 and 500 bpm, respectively, while no strains fell above the population reference range. The PR interval normal reference range is 20.14 – 32.75 ms. While no strains are below the population reference range, CC012, CC044, and CC057 are above at 35.9, 30.4, and 34.6 ms, respectively. The normal reference range for QRS complex is 8.45 - 14.15 ms. Only CC057 and is above the reference range at 17.5 ms. While CC026 still falls in the population range at 8.78, this strain had the shortest QRS complex. QTc normal reference range is 40.48 – 52.26 ms. No strains are below this range, but CC011, CC012, and CC057 are above the reference range at 52.6, 54.3 and 56.7 ms, respectively.

### Mouse models of extreme cardiac phenotypes

The differences in cardiac parameters amongst the strains may be exploited to identify natural models for various disease-like cardiac phenotypes. Based on the echocardiography findings, several CC strains show disease-like phenotypes observed in humans (Table 1). From the anesthetized echocardiography data, CC018 has EDV within the normal range but higher ESV and low EF, hallmarks of dilated cardiomyopathy. CC023 and CC029 have relatively normal EF but low GLS and FS (CC023), which can be seen in patients with early signs of cancer therapy-related cardiac dysfunction. CC031 has dilated EDV, and increased SV and CO, which is associated with high-output heart failure such as with chronic amemia or hyperthyroidism. From the conscious echocardiography data, CC010 has dilated EDV, increased SV and decreased HR, which are often seen in “athlete’s hearts” but can also be seen in dilated cardiomyopathy during compensation. CC060 has high ESV but low EF and FS, consistent with systolic heart failure or heart failure with reduced ejection fraction (HFrEF) that can occur in chronic hypertension or valvular heart disease among other conditions. CC035 has high ESV and low HR that can occur in bradycardia-related dilated cardiomyothaphy.

From the electrocardiogram data, potential new models of long QT syndrome were identified. CC057 has low HR with high PR, ST, QRS, and QTc intervals suggesting condution system abnormalities such as those associated with ischemica heart disease or fibrosis of the conduction system. Similarly, CC012 has low HR with high PQ, PR, ST, and QTc that is also suggested of a conduction system abnormality with delayed ventricular repolarization that can be caused by hypothyroidism.

## Discussion

In the present study, cardiac physiology was evaluated in 58 strains of the CC and C57BL/6J, a classical inbred strain that is widely used as a single-strain reference. Overall, our findings demonstrate that there are marked differences in cardiac physiology and response to anesthesia among strains, and that these differences are due to genetic background. While including genetic diversity adds to the scale of the study, defining what is “normal” for mouse cardiac function can better inform selection of strains as models as has been successfully done previously for cancer (DORMAN *et al*. 2016; WANG *et al*. 2019), toxicology (CICHOCKI *et al*. 2017; LEWIS *et al*. 2019; ZEISS *et al*. 2019), immunology (GRAHAM *et al*. 2017; LORE *et al*. 2020), neuroscience (MOLENHUIS *et al*. 2018), and reproduction (SHORTER *et al*. 2019).

The first key finding of this study was the large amount of variation in cardiac phenotypes under normal echocardiography conditions. All traits monitored in this study were highly heritable. Anesthetized echocardiography was slightly more heritable than conscious echocardiography, likely due to anesthesia minimizing some uncharacterized sources of variation. This naturally occurring variation more closely models that seen in the human population, redefining what “normal” means for mouse models and method of echocardiography (CHOQUET *et al*. 2020). The anesthetized volume parameters %CV are comparable to those seen in humans, while conversely the conscious functional parameter, EF, is more similar to humans. This indicates that primary measurements are perhaps more accurately measured under anesthesia in mice. However the effect of anesthesia can drastically change functional parameters, and those values should be recorded when the mouse is conscious.

The second finding was that anesthesia and stress of conscious mouse handling can differentially influence echocardiographic phenotypes based on genetic background. While residual effects contribute to a large portion of the overall variability seen across strains in EF, GLS, and especially HR, the strain effects are also highly significant and account for the largest source of variation. Residual influence is smaller in anesthetized mice, which supports that conscious echocardiography is inherently noisier and likely due to experimental variables such as the level of stress, physical scruffing, and background noise. EF has been the most common and established method for assessment of LV function in humans, but in recent years GLS has shown to be more sensitive to LV dysfunction at earlier diagnostic stages (POTTER AND MARWICK 2018), and has improved reproducibility (KARLSEN *et al*. 2019) and precision (POTTER AND MARWICK 2018). The American Society of Echocardiography and the European Association of Cardiovascular Imaging have agreed that deformity changes precede ventricular dysfunction, and GLS has become a routine diagnostic for cardiotoxicity prediction in cancer patients (GRIPP *et al*. 2018). In normal and dilated human hearts, EF is related to GLS by a factor of approximately 3 (ONISHI *et al*. 2015). However, in human diseases where the ratio of SV to LV cavity size can be preserved, such as in chronic kidney disease (KRISHNASAMY *et al*. 2015) or recently emerging heart failure with preserved EF, EF is not informative and is not related to GLS (ANDERSSON AND VASAN 2014).

It is noteworthy that the differential response to anesthesia, which affects EF or GLS measurements or both, is also evident in the data from this study. CC035 and CC060 have little change in EF and GLS between anesthesia and conscious echocardiography, whereas B6, CC010, CC018, CC027, and CC045 are drastically different by a factor of more than 2.5. Strains with a lower EF under anesthesia seem to maintain the highest intrastrain variability, indicating that a high and healthy baseline EF is more reproducible. In animals with preexisting cardiac pathology or experimentally-induced cardiac pathology, reduced myocardial contractility and autonomic nervous system suppression via anesthesia can exaggerate cardiac pathologies and confound interpretation (YANG *et al*. 1999). Of note is the dramatically lower EF under anesthesia in a few strains; while most experts would consider this heart failure, the method of anesthesia was the same for each individual. Mice were breathing normally and recovered normally, even at these values. While strain is a factor influencing variability, higher H^2^ values of phenotypes when conscious indicate that anesthesia reduces residual effects that have a genetic contribution. Under both conscious and anesthetized echocardiography, outlier strains outside the established reference ranges are potential models of human cardiac disease. Some strains that are models of, for example, dilated cardiomyopathy under anesthesia are not the same strains that present as models under conscious echocardiography. This indicates that not only does genetic background need to be taken into account when defining models for human cardiac disease, but the method of echocardiography does as well. The outlying strains listed as potential models for various cardiac disease states are a useful tool for researchers looking for naturally pre-disposed models of cardiac phenotypes, in both conscious and anesthetized echocardiography.

The third key finding was that genetic background influences cardiac electrophysiology, and the redefinition of normal cardiac electrophysiological intervals in mice. Residual effects have a large contribution to the variability seen across strains, but strain differences still contribute significantly. It is still a matter of debate whether the mouse is an acceptable model for studies of human cardiac electrophysiology (FARRAJ, HAZARI AND CASCIO 2011; BOUKENS *et al*. 2014). The extrapolation to humans of the T wave, methods that induce diseases such as myocardial ischemia and Brugada syndrome (BOUKENS *et al*. 2014), and overall cardiac morphology are a few reasons why this must be done with caution. Mice are typically not used for their electrophysiological traits in drug safety evaluation, but are used in studies of chemical toxicity, and therefore still need to be characterized and optimized to be the most accurate model. Under anesthesia, the B6 mouse appears to be an average mouse as this strain was among the strains with highest conscious EF, and yet under anesthesia was a more intermediate strain at 14^th^ lowest. Heritability of HR during ECG was high (H^2^ = 0.44), although not as high as other phenotypes, indicating environmental factors, and therefore sympathetic tone, play a large role in maintaining beat rate. Other significant ECG phenotypic variances were generally smaller within than between strains, indicative of elevated trait heritabilities. The B6 mouse is used most commonly when comparing mouse electrophysiology to humans, especially in risk assessment research. The duration of action potential, as indicated by QTc, is an important interval for both congenital and induced pathologies, especially for cardiotoxicity. Although the B6 mouse does fall within the reference range, it is remarkable that this strain has among the shortest QTc of the 59 mouse strains analyzed, and therefore even less accurately represents the “normal” human. This raises concerns for this strain’s abundant use in mouse cardiac studies and general representation of human heart health.

The fourth key finding is the amount that environment affects HR, even in a diverse population, indicating that an average HR is relative and cannot be reduced to one value. Sex contributes significantly to variability of HR difference between anesthetized and conscious echocardiography, indicating that male and female mice respond differently to stressors of conscious and anesthetized echocardiography. HRs in anesthetized echocardiography, conscious echocardiography, and ECG were significantly different (p < 0.0001), although the reference range determined using conscious echocardiography and ECG were similar. While some strains maintained their outlier state among strain ranking for HR values, others had more scattered rankings. HRs were among the least heritable traits, and yet the change in HR from conscious to anesthetized mice was heritable, indicating that the response of HR to anesthesia is heritable. Strains like CC005 have one of the lowest HRs under anesthesia, and yet maintain intermediate level EF values, further demonstrating that HR is not necessarily the constant variable by which to evaluate anesthetized echocardiography phenotypes. Consistent with these observation, some humans can have resting HR that are up to 70 bpm different from other individuals (QUER *et al*. 2020).

The fifth key finding is the identification of new, natural models of human cardiac disease predisposition. There is a pressing need to understand the molecular mechanisms of cardiovascular disease, and understanding even baseline genetic differences can have implications for response to toxicants, drugs, and environmental exposures. Indeed, one study found differential response to doxorubicin-induced cardiotoxitiy (ZEISS *et al*. 2019), and yet the susceptible strains still do not correlate with the current study’s at risk strains. Baseline phenotypes might not be directly informative for predicting outcomes of drugs or chemicals, but to elucidate the molecular mechanisms underlying cardiac pathologies the depth of baseline diversity must first be explored. The current study’s models have advantages compared to cardiac models prepared with genetically engineered mice, xenobiotic-induced remodeling, or surgical intervention. Transgenic mice are complicated and expensive to engineer; manipulating the mouse at the genetic level will not always develop into a disease that is comparable to humans. Genetic engineering is not entirely foolproof and sometimes the work needs to be repeated to obtain the desired result.

Unwanted side-effects from drug or xenobiotic cannot be avoided, and established doses may underestimate or overestimate doses needed to produce the desired result in a diverse population. Surgical models of cardiac disease involve rapid induction of the stressor, whereas the current study’s models develop more naturally over the course of the mouse’s life, similar to humans.

While the current study has strengths in terms of the number of strains, time of data collection, and the use of standard echocardiographic techniques; several limitations must be acknowledged. Intrastrain variance in some strains was high with some strains being more differentially susceptible to external stressors, indicating that reproducibility is in itself a phenotype that must be acknowledged. This can also be observed in inter-laboratory differences contributing to variability. We did not observe any sex differences in the functional phenotypes, most likely due to the low number of replicas per strain. The echocardiographic and electrocardiographic software analysis were designed based on data from only a few inbred strains, which could lead to inaccuracies in analysis when the software assumes similar heart tissue density and shape across all animals. To conclude, this study reports the largest cardiac phenotyping analysis performed in a genetically diverse mouse reference population that has supported the development of cardiac reference ranges for *Mus musculus* and identification of new mouse models for cardiac disease predisposition.

## Data Availability Statement

The data underlying this article are available in The Mouse Phenome Database at https://phenome.jax.org/

## Acknowledgements

The authors thank Dr. Nadia Rosenthal from The Jackson Laboratory for sharing raw data from Salimova et al. and The Jackson Laboratory for posting raw data on the Mouse Phenome Database.

